# Signal peptidase complexes set species-specific rules for signal peptide cleavage

**DOI:** 10.64898/2026.09.20.752496

**Authors:** Yeonji Chung, Gilberto P. Pereira, Young June Lee, Achille Agnesa, Lisbeth R. Kjølbye, Henrik Nielsen, Gunnar von Heijne, Paulo C. T. Souza, Hyun Kim

**Affiliations:** School of Biological Sciences and Institute of Biodiversity, Seoul National University, Seoul 08826, South Korea; Laboratoire de Biologie et Modélisation de la Cellule, UMR 5086, CNRS, Ecole Normale Supérieure de Lyon, 46 Allée d’Italie, 69364, Lyon, France; Centre Blaise Pascal de Simulation et de Modélisation Numérique, Ecole Normale Supérieure de Lyon, 47 Allée d’Italie, 69364 Lyon, France; Zymvol Biomodeling, Barcelona, Spain; Bioengineering Department, University Cote d’Azur, France; Molecular Microbiology and Structural Biochemistry, CNRS UMR 5086 and Université Claude Bernard Lyon 1, Lyon, France; Section for Bioinformatics, Department of Health Technology, Technical University of Denmark, Kongens Lyngby, Denmark; Department of Biochemistry and Biophysics, Stockholm University, SE-106 91 Stockholm, Sweden; Science for Life Laboratory Stockholm University, Box 1031, SE-171 21 Solna, Sweden; School of Dentistry, Seoul National University, Seoul 08826, South Korea

**Keywords:** Signal sequence, signal peptide, signal-anchor, secretion, membrane protein

## Abstract

Protein secretion is essential for cell function and begins when signal peptides direct nascent proteins into the secretory pathway, where they are cleaved by the signal peptidase complex (SPC). Although this pathway is highly conserved, signal peptides do not always function efficiently across species, and the molecular basis for this incompatibility remains unclear. Here, we show that human signal peptides with longer hydrophobic cores efficiently target and translocate nascent proteins in both yeast and human but are frequently left uncleaved in yeast, identifying signal peptide cleavage as a major barrier to cross-species compatibility. Molecular dynamics (MD) simulations reveal that, despite their highly conserved architectures, yeast and human SPCs generate distinct membrane-thinning profiles near the signal peptide-binding region. Our results support a model in which SPC–lipid interactions tune the local membrane environment to accommodate signal peptides of distinct hydrophobic core length, providing a biophysical mechanism for species-specific signal peptide recognition. These findings offer a mechanistic framework for understanding and potentially engineering protein secretion across diverse expression hosts.

## Introduction

Protein secretion is an essential process for cell growth and survival. Secretory proteins are first targeted to the plasma membrane in prokaryotes or to the endoplasmic reticulum (ER) membrane in eukaryotes through N-terminal signal sequences. Signal sequences direct the early biogenesis of secretory proteins through three sequential steps. First, signal sequences mediate targeting of nascent chains to the Sec translocon or the ER membrane protein complex (EMC) in the membrane (1–3). Second, engagement of the signal sequence with the Sec translocon initiates translocation and membrane insertion of the nascent polypeptide (4–7). Finally, the signal sequence is cleaved by signal peptidase on the periplasmic side of the bacterial membrane or within the ER lumen in eukaryotes, allowing the nascent chain to fold into its mature conformation (8).

Signal sequences possess a tripartite organization consisting of an N-terminal (n) region, a central hydrophobic (h) region, and a C-terminal (c) region containing the signal peptidase cleavage site (9). Positively charged residues are enriched in the n-region, whereas the h-region is composed predominantly of hydrophobic, helix-forming amino acids. The cleavage site is defined by the presence of small, neutral amino acids at the -1 and -3 positions relative to the cleavage site (10, 11). Despite these conserved features, signal sequences exhibit remarkable diversity in both length and amino acid composition, with little overall sequence conservation (12–14). The mechanisms governing SRP recognition and cotranslational targeting have been extensively characterized, yet considerably less is known about how signal peptidases distinguish their substrates.

The catalytic domains of *Escherichia coli* leader peptidase (LepB) and the human signal peptidase complex (SPC) share a conserved catalytic structure that includes a substrate-binding groove (15–17). However, their overall architectures differ substantially. Whereas bacterial signal peptidase functions as a monomer, the eukaryotic SPC is a hetero-tetrameric complex consisting of the catalytic subunit SEC11A or its paralog SEC11C together with three additional subunits (SPCS1-3). Cryo-EM structures of the human SPC revealed a unique architecture in which the transmembrane (TM) helices of the SEC11A/C-SPCS2 pair and the SPCS3-SPCS1 pair form two separate bundles, creating a membrane-embedded gap referred to as the TM window (16). A recent structure captured the h-region of a signal peptide bound within this TM window (17). Molecular dynamics (MD) simulations further demonstrated that the lipid bilayer within the TM window is thinner than the surrounding membrane (16, 18). Together, these structural observations suggest that membrane thinning within the TM window serves as a molecular ruler that preferentially accommodates signal peptides with short h-regions while discriminating against signal-anchor sequences.

Because both the secretory machinery and the tripartite architecture of signal peptides are evolutionarily conserved, many signal peptides function across species (19–21). Consistent with this conservation, numerous rat signal peptides have been identified based on their ability to direct secretion of a reporter protein in *S. cerevisiae* (22). However, not all mammalian signal peptides function efficiently in yeast (23–25). Conversely, the signal peptide of yeast carboxypeptidase Y (CPY) is inefficient in mammalian cells unless its hydrophobicity is increased (24, 26). In yeast, signal peptides with relatively weak hydrophobicity or shorter h-regions are often not recognized by the signal recognition particle (SRP) and instead utilize the post-translational Sec62/Sec63-dependent translocation pathway (27–30). In both yeast and mammals, the hydrophobicity of signal peptides is a major determinant of SRP recognition and Sec translocon engagement. However, whether signal peptide cleavage is compatible between species, and which features govern efficient cleavage by the SPC in different organisms, remain largely unknown.

To address these questions, we have now investigated the compatibility of human signal peptide cleavage in *S. cerevisiae*. Using model substrates containing poly-leucine h-regions of varying lengths, we determined the maximal h-region length threshold required for efficient signal peptide cleavage in yeast and human cells. The 50% cleavage threshold was approximately three residues shorter in yeast than in human cells. We next compared the processing of natural human signal peptides fused to a CPY reporter in both organisms. Although most signal peptides with relatively long h-regions were efficiently targeted and translocated in both yeast and human cells, the majority were efficiently cleaved only in human cells and remained largely uncleaved in yeast. Consistent with these observations, comparative analysis revealed that *S. cerevisiae* signal peptides are generally shorter than *Homo sapiens* signal peptides. Finally, MD simulations of yeast and human SPC embedded in the lipid bilayer revealed marked differences in membrane thinning within the TM window.

Collectively, our results establish membrane thinning within the SPC TM window as a molecular ruler that determines species-specific h-region length thresholds for signal peptide cleavage. These findings provide new mechanistic insights into how lipid interaction with SPC defines species-specific substrate selectivity during signal peptide processing.

## Results

### The h-region length threshold for signal sequence cleavage differs between yeast and human SPC

Previous studies using poly-leucine (Leu) model signal sequences showed slightly different cleavage patterns between cell-free dog pancreatic microsomes and *in vivo* yeast cells (31, 32).

To determine if these differences are inherent to the species, we compared signal sequence cleavage in *S. cerevisiae* and human HEK293T cells using a set of LepCC variants (31, 32) (Fig. 1A). These model proteins contain an N-linked glycosylation site to monitor ER targeting and a C-terminal hemagglutinin (HA) tag for detection by Western blotting (Fig. 1A). LepCC variants were glycosylated in both cell types, indicating their ER targeting (Fig. 1B and Fig. S1). As a control for the SPC-uncleaved form of LepCC variants, a cleavage site mutant where the +1 position is changed to Proline (LepCC (14L+P1)) was used (33–35). In yeast cells, efficient SPC-mediated cleavage occurred for signal sequences with 15 or fewer Leucines in the h-region (Fig. 1B and C). In comparison, in human cells, over 50% cleavage was maintained for signal sequences containing up to 18 Leu residues in the h-region (Fig. 1B and C).

**Figure 1.**
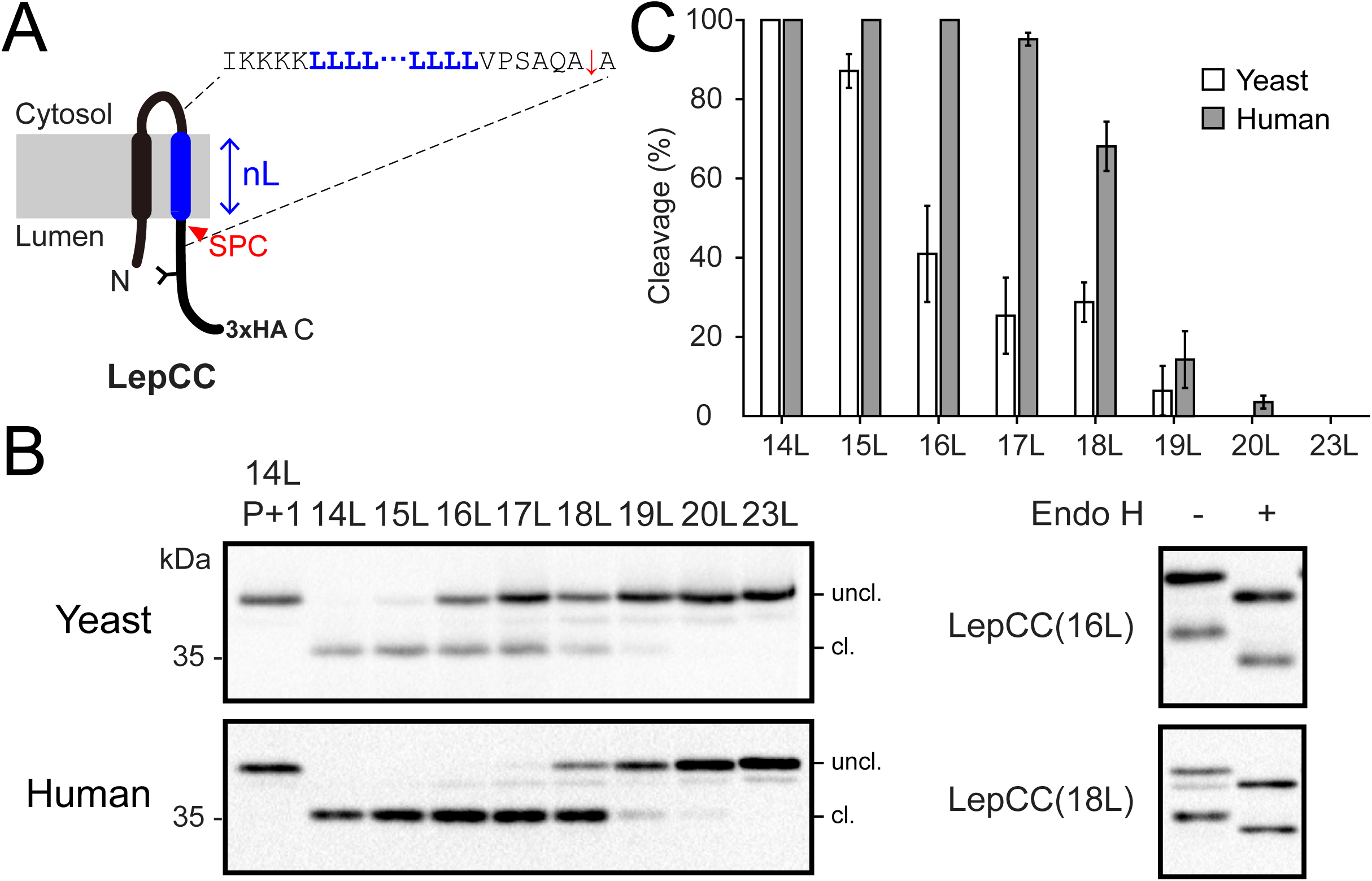
The h-region length threshold for signal sequence cleavage in yeast and human cells. (A) Schematic of LepCC variants. nL indicates the number of leucine residues (n=14 to 23). Y indicates an N-glycosylation site. nL and flanking sequences are shown. Red arrow indicates SPC cleavage site. (B) Western blot analysis of cleavage of LepCC variants in yeast (W303-1α) and human cells (HEK293). LepCC(14L) P+1 is a cleavage site mutant where a residue in position +1 relative to the cleavage site was substituted with a proline. Endoglycosidase H (Endo H)-treated LepCC(16L) from yeast and LepCC(18L) from human cells are shown on the right. (C) Quantifications of western blot data. Error bars indicate mean± standard deviations (S.D).

These results indicate that the 50% cleavage threshold for the h-region length of signal sequences is approximately 16 residues for yeast SPC, and 18-19 residues for human SPC. This reveals a distinct three-residue species difference in the h-region length of signal sequences that are processed by yeast and human SPC.

### Human signal peptides with longer h-region are uncleaved by yeast SPC

To validate the species-specific 50% cleavage threshold observed with model signal sequences, we assessed the processing of selected human signal peptides with predicted h-regions longer than 16 residues in yeast cells. We hypothesized that these human sequences would be resistant to cleavage by the yeast SPC.

We utilized a truncated carboxypeptidase Y (CPY) fusion reporter, which contains three N-linked glycosylation sites to monitor ER targeting (Fig. 2A). The N-terminal 60 amino acids of the selected human secretory proteins were fused to the CPY reporter (Fig. 2A). To ensure comparability with our LepCC model variants, we selected human SPs with predicted h-regions longer than 16 residues, an h-region hydrophobicity corresponding to ΔG < -1 kcal/mol (36) and an n-region longer than 5 residues (Table 1).

**Figure 2.**
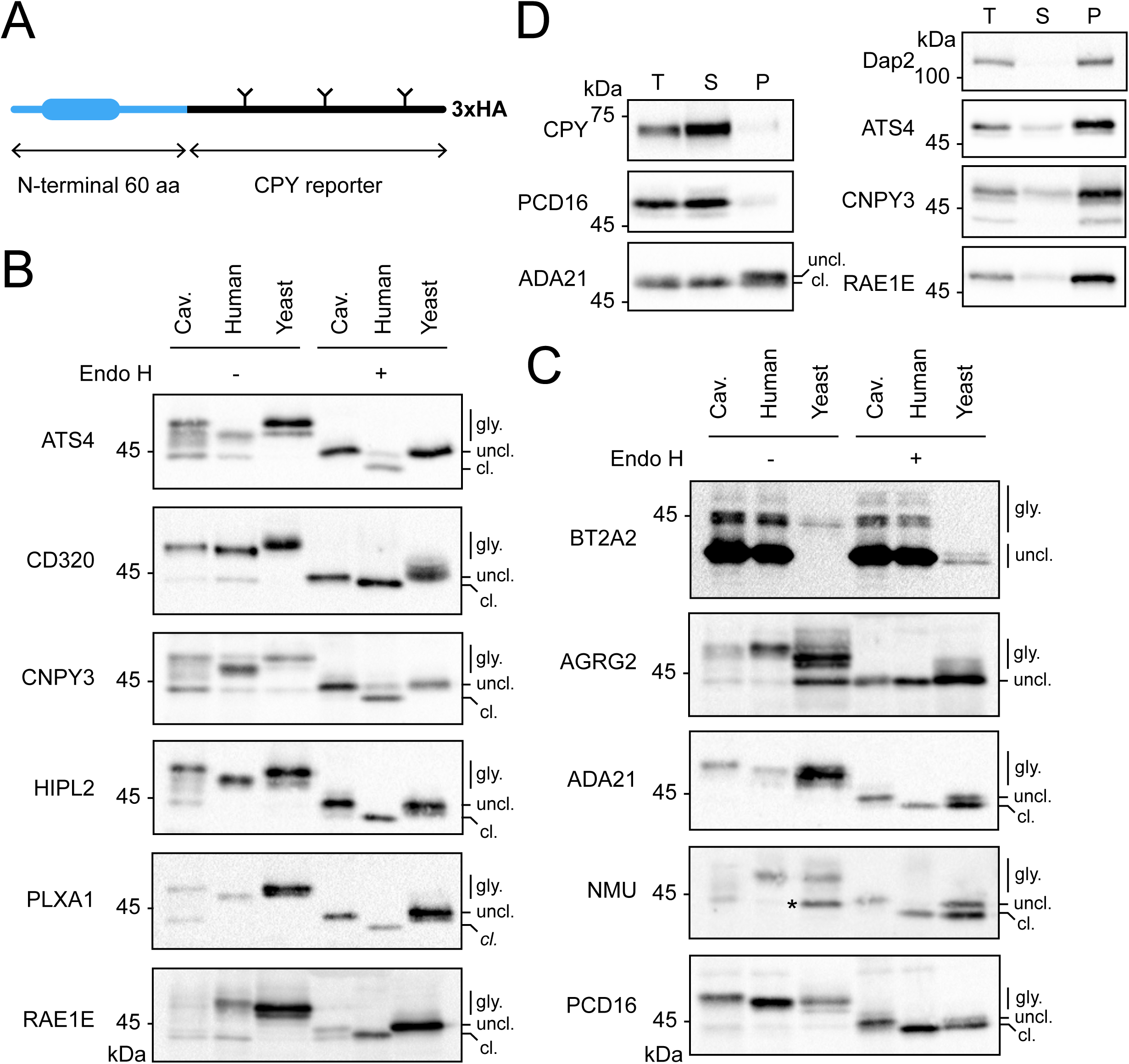
SPC-mediated cleavage of human signal peptides in yeast and human cells. (A) Schematic of human signal peptide fused to a truncated CPY reporter. Y indicates an N-linked glycosylation site. (B) and (C) Western blot analysis of fusion proteins of human signal peptide. Cav is cavinafungin-treated HEK293 cells used as a control for uncleaved signal peptide. (D) Carbonate extraction of ATS4-CPYt, CNPY3-CPYt and RAE1E-CPYt expressed in yeast cells. CPY and Dap2 are soluble and membrane protein controls, respectively. T, total, S, soluble, P, pellet fractions of carbonate extraction.

**Table 1.**
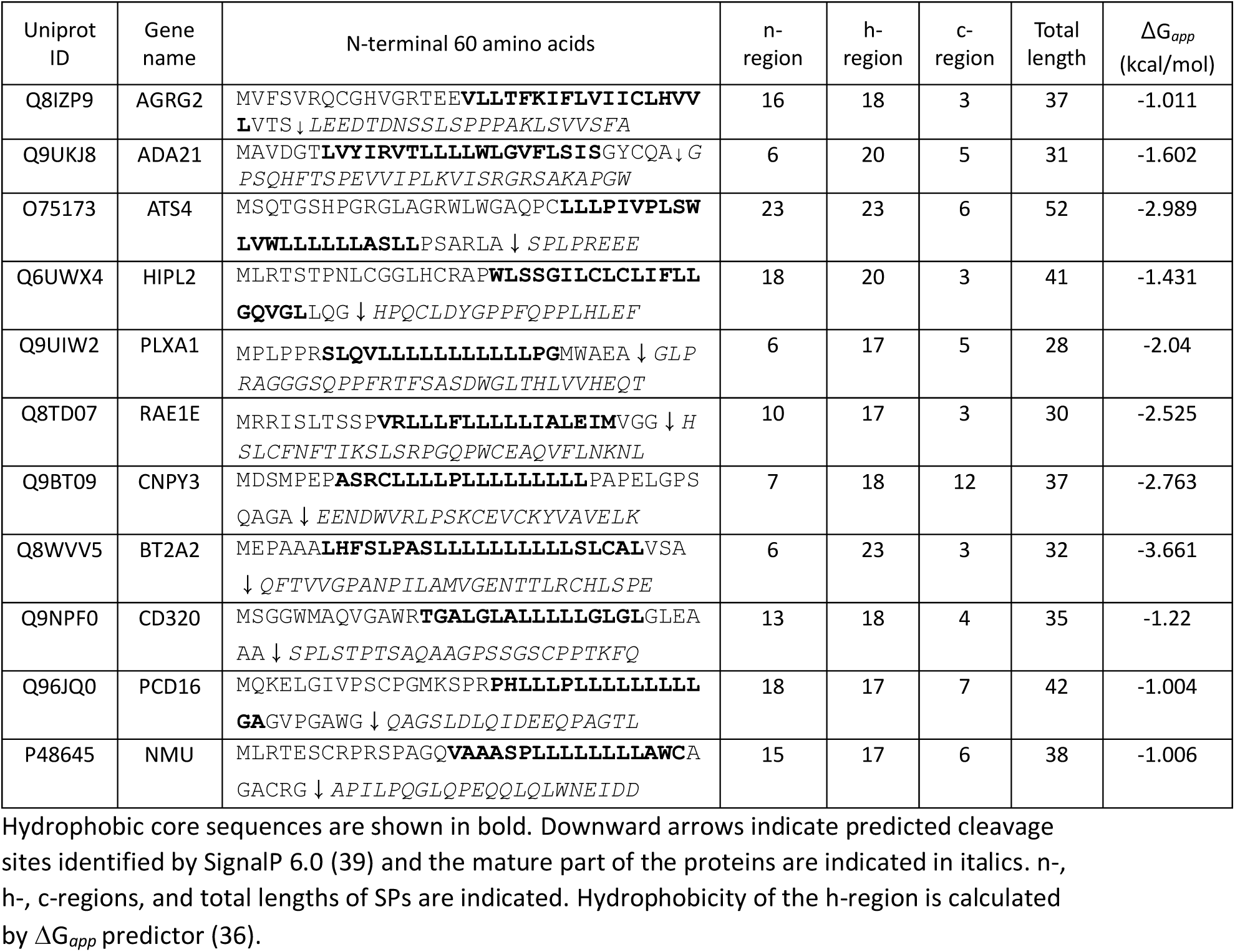
List of human signal peptides (SPs) used in this study.

Of the eleven fusion proteins tested, all were glycosylated in yeast cells, indicating that these human signal peptides mediate proper targeting and translocation in yeast (Fig. 2B-C). In HEK293T cells, all were glycosylated, but BT2A2 was only partially glycosylated (Fig. 2C). Glycosylated BT2A2 in human cells was endoglycosidase H (Endo H) resistant and peptide:N-glycosidase (PNGase) sensitive, indicating that it was trafficked through the Golgi apparatus (Fig. S1B). To distinguish signal sequence cleaved and uncleaved forms, HEK293T cells were treated with cavinafungin (Cav), an inhibitor of the eukaryotic SPC (37). The bands in Cav-treated samples thus indicate signal sequence uncleaved form. AGRG2 remained uncleaved in both human and yeast cells, and targeting in yeast was inefficient (Fig. 2C). AGRG2 belongs to the adhesion GPCR family, which produces diverse mRNA transcript variants that can subsequently give rise to multiple isoforms (38). Interestingly, some of these transcripts have been shown to encode proteins with eight TM domains as a result of a signal anchor sequence instead of a signal peptide at the N-terminus (38). AGRG2 is predicted to have a borderline signal peptide by SignalP 6.0 (39), AlphaFold predictions, and is predicted by some membrane protein prediction algorithms to yield an eight-TM protein (Fig. S2). These predictions and our data suggest that the N-terminal signal sequence of AGRG2 may function as a signal anchor sequence rather than a cleavable signal peptide in human cells. For six of the fusion proteins, the signal sequence remained uncleaved in yeast cells but efficiently processed in human cells (Fig. 2B). These data confirm that human signal peptides with longer h-regions are recognized and cleaved by the human SPC but are inefficiently cleaved by the yeast SPC, consistent with our findings using model signal sequences in Fig.1.

To determine the membrane association of cleaved and uncleaved human signal peptides in yeast cells, we performed carbonate extraction on five fusion proteins (PCD16, ADA21, ATS4, CNPY3 and RAE1E) along with control soluble and signal-anchor proteins, CPY and Dap2. Upon carbonate extraction, the PCD16 fusion protein was mostly detected in the soluble fraction, whereas the ATS4, CNPY3 and RAE1E fusion proteins were detected in the pellet fraction, indicating that the uncleaved human signal peptides remained anchored in the membrane (Fig. 2C). For the ADA21 fusion protein, the slowly-migrating species containing uncleaved signal peptide was detected in the pellet fraction, whereas the faster-migrating species containing cleaved signal peptide was detected in the soluble fraction. These data demonstrate that some human signal peptides containing longer h-regions remain uncleaved in yeast SPC and become signal-anchor sequences.

### The tripartite features of natural signal sequences are less distinct compared to model signal sequences

For three of the tested fusion proteins, SPC-mediated cleavage was observed in both human and yeast cells (Fig. 2C). Unlike the model LepCC signal sequences, which possess well-defined n-, h-, and c-regions (Fig. S3A), the boundaries of these regions are less distinct in natural signal sequences (Fig. S3B). Broad overlaps between these regions indicate that an actual h-region might be shorter than predicted, and be below the cleavage threshold for yeast SPC. To test this, we lengthened the h-regions of those three natural signal sequences by adding two Leucines (+2L) and assessed their processing in human and yeast cells.

The addition of two Leucines to the h-region of the ADA21 signal sequence did not affect cleavage in human cells, but rendered the protein mostly uncleavable in yeast (Fig. 3A). For NMU, initial ER targeting was inefficient in yeast, as approximately half of the protein remained unglycosylated (Fig. 2C, Fig. 3B, the 3^rd^ lane). Adding +2L to the NMU h-region markedly reduced the amount of unglycosylated protein, indicating improved ER targeting (Fig. 3B). While this modification did not affect cleavage in human cells, the amount of SPC-uncleaved protein markedly increased in yeast (Fig. 3B).

**Figure 3.**
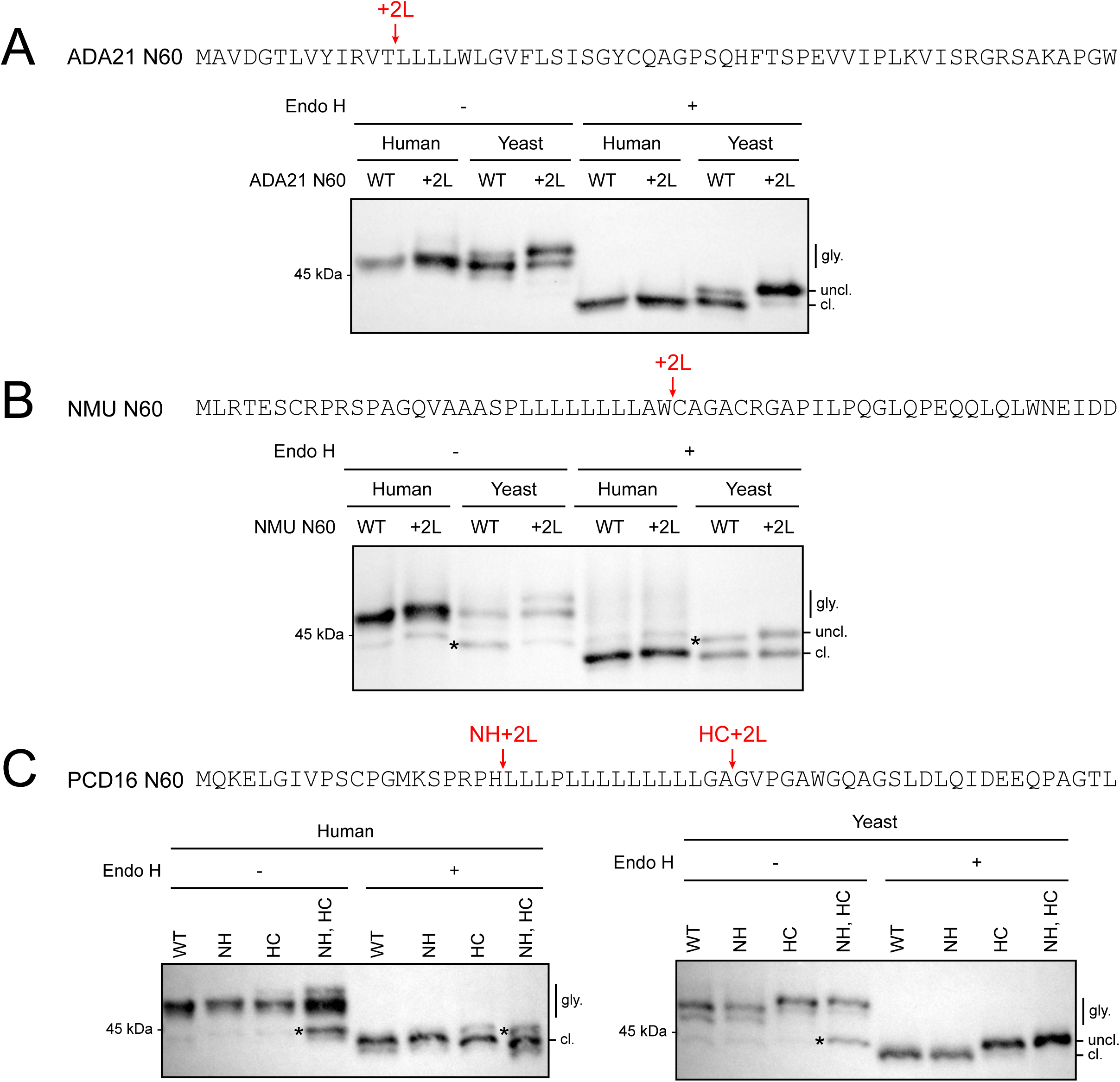
SPC-mediated cleavage of human signal peptides with extended h-region in yeast and human cells. (A) N-terminal 60 aa sequence of ADA21; Red arrow indicates the position where two leucines were added (+2L) (top) and western blot analysis of ADA21 and ADA21 +2L (bottom). (B) N-terminal 60 aa sequence of NMU; Red arrow indicates the position where two leucines were added (+2L) (top) and western blot analysis of NMU and NMU +2L (bottom). (C) N-terminal 60 aa sequence of PCD16; Red arrows indicate the positions where two leucines were added (+2L) either at n/h-region junction or h/c-region junction (NH+2L and HC+2L, respectively), and at both n/h-region and h/c-region junctions (NH, HC +2L) (top) and western blot analysis of PCD16, PCD16 NH+2L, PCD16 HC+2L and PCD16 NH, HC+2L (bottom). * indicates untargeted proteins.

Consistently, addition of +2L at the PCD16 h/c-region junction (HC+2L) mostly did not affect cleavage in human cells but prevented cleavage in yeast. However, the effect was position-dependent as adding +2L to the n/h-region junction (NH+2L) had no effect on cleavage either in human cells or yeast (Fig. 3C, NH +2L). This could be due to the presence of multiple potential cleavage sites. In addition to the SignalP 6.0 prediction, we thus conducted a SignalP 4.1 (40) analysis of the PCD16 sequence, since SignalP 4.1 better than 6.0 indicates alternative cleavage sites. Although SignalP 4.1 (40) predicts residues 40-42 (AWG) as the primary cleavage site (Fig. S3C), residues 34-36 (GAG) and 37-39 (VPG) are both potential SPC cleavage sites as SPC cleaves after small, neutral amino acids at -3 and -1 positions (40). Given that addition of +2L between residues 35A and 36G abolished cleavage in yeast cells, it may be that the 34-36 (GAG) is used in yeast cells whereas downstream sites are used in human cells. Interestingly, adding Leucines to both ends of the h-region mildly impaired ER targeting in both cell types (Fig. 3C), likely because the h-region exceeded the 15-16 residue optimal length for SRP binding, therefore reducing ER targeting efficiency (41). Despite mild defect in targeting, proteins that reached the ER were still cleaved by SPC in human cells but not in yeast (Fig. 3C). Collectively, these data further support the conclusion that signal peptides with longer h-regions are processed by human SPC but are excluded by the yeast complex.

### Some yeast signal-anchored proteins are cleaved by human SPC

As some human signal peptides become signal-anchor sequences when expressed in yeast, we wondered whether yeast signal-anchored proteins could be cleaved by SPC when expressed in human cells. To test this idea, we selected seven yeast signal-anchored proteins that contain potential SPC cleavage sites (Fig. S4) and subcloned the genes into mammalian expression vectors. We added two N-linked glycosylation sites (2g) for those lacking endogenous N-linked glycosylation sites to check their ER targeting, along with triple HA tag at the C-terminus for detection by western blotting, and assessed their SPC-mediated cleavage in human cells (Fig. 4). Mnn11, Atg15 and Hoc1 were truncated after amino acid 350 and fused with HA tag to make easier to distinguish signal sequence cleaved and uncleaved proteins on a SDS gel by size differences (denoted as t).

**Figure 4.**
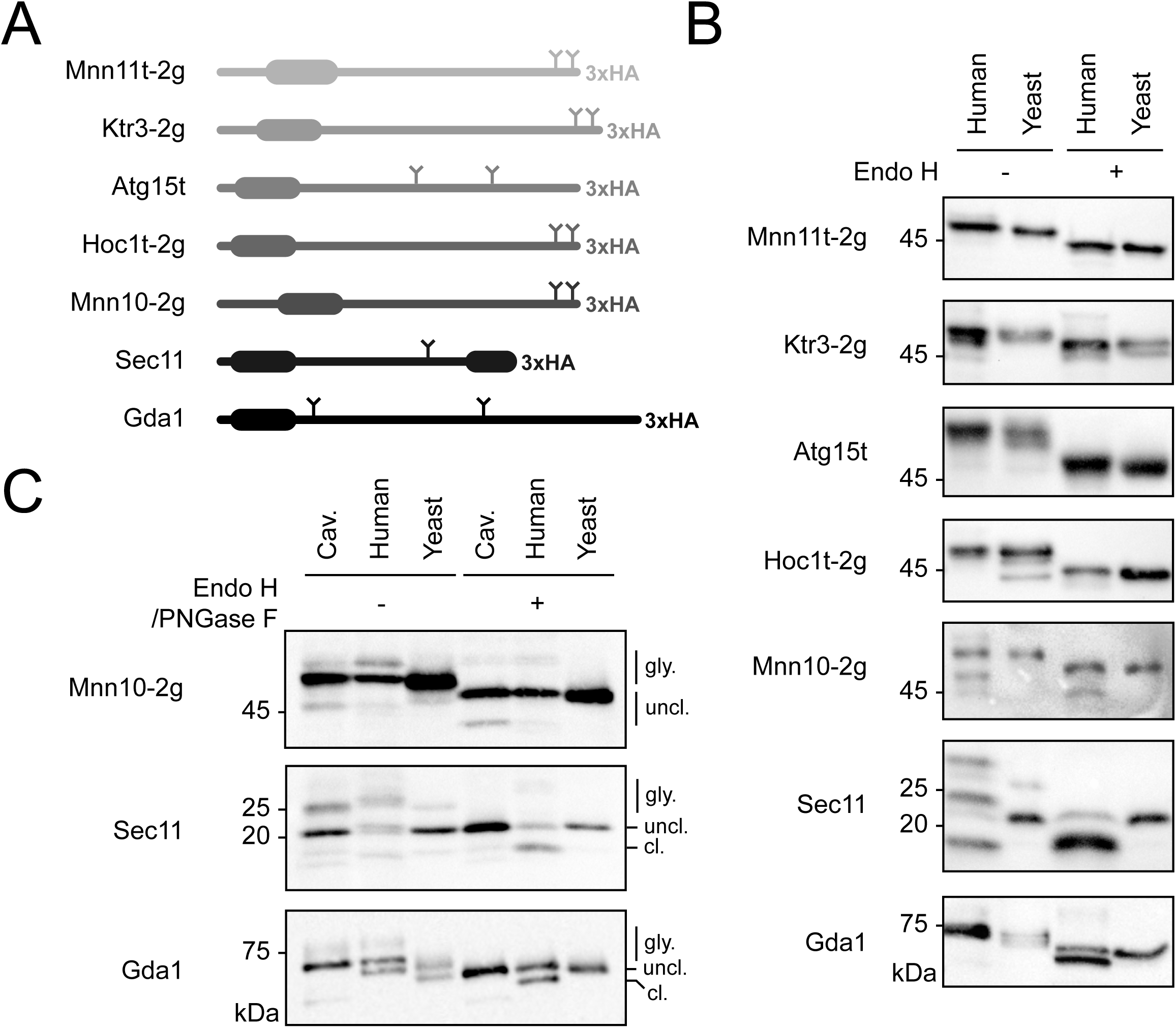
SPC-mediated cleavage of yeast signal-anchored proteins in yeast and human cells. (A) Schematics of 7 yeast signal-anchored proteins. 2g refers to two N-linked glycosylated sites added at the C-terminal end of Mnn11t, Ktr3, Hoc1t and Mnn10. (B) and (C) Western blot analysis of yeast signal-anchor proteins expressed in HEK293 cells treated with or without Cav, and yeast cells. Samples were treated with Endo H or Peptide:N-glycosidase F (PNGase F).

All seven proteins were glycosylated in human cells, indicating that they were properly targeted to the ER. Sec11 has an endogenous, inefficiently modified N-linked glycosylation site (42). Comparing the size of the proteins expressed in yeast and human cells, three of them showed smaller size in human cell lysates compared to yeast cell lysates (Fig. 4B). To confirm that those smaller size proteins in human cell lysates are SPC-cleaved products, we assessed protein expressions of Mnn10-2g, Sec11 and Gda1 from human cells treated with and without SPC inhibitor, Cav (Fig. 4C). For Sec11 and Gda1, smaller size proteins were no longer detected in human cells treated with Cav, indicating that they are indeed SPC-cleaved proteins. For Mnn10-2g, a faint smaller size band remained in the Cav-treated human cells, meaning that this form is not generated by SPC-mediated cleavage. These data show that some yeast signal-anchor sequences are cleaved by human SPC.

### Comparison of yeast and human signal peptide length

Our experimental data show that signal sequences having longer h-regions are efficiently cleaved by human but not by yeast SPC, implicating that human signal peptides can possess longer h-regions than those of yeast. To determine if this reflects a proteome-wide trend, we compared signal peptides from yeast and human databases (https://services.healthtech.dtu.dk/services/SPEx-1.0/)(Table S1). While the median and mean overall lengths of human signal peptides are two residues longer than those of *S. cerevisiae* (Fig. 5A), this difference is primarily driven by longer n-regions in the human sequences (Fig. 5B).

**Figure 5.**
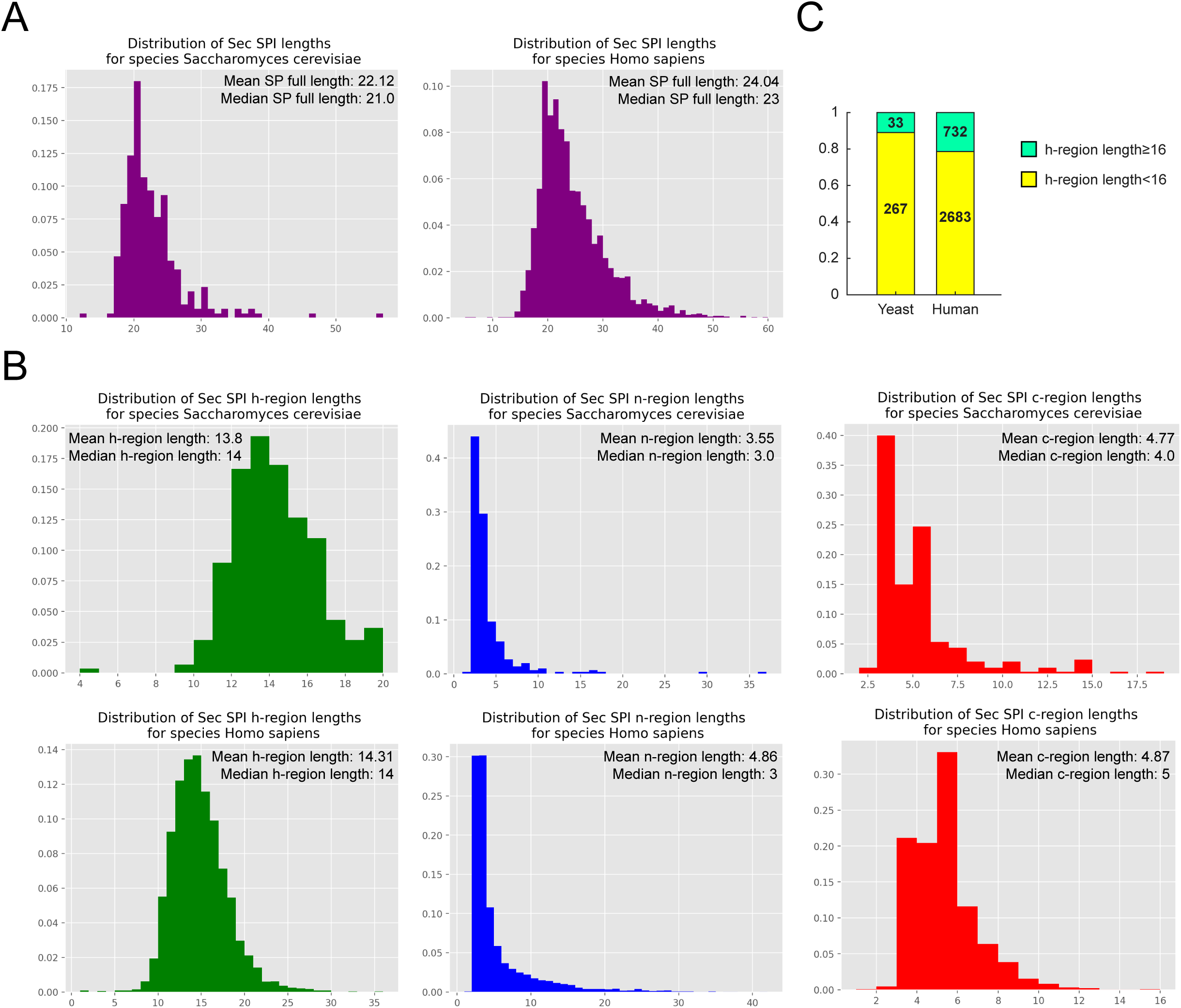
Comparison of yeast and human signal peptide length. (A) Distribution of total lengths of signal sequences in yeast (left) and humans (right) (B) Left: Distribution of h-region lengths of signal sequences in yeast (top) and humans (bottom). Middle: Distribution of n-region lengths of signal sequences in yeast (top) and humans (bottom). Right: Distribution of c-region lengths of signal sequences in yeast (top) and humans (bottom). (C) Bar graph showing the proportion of signal sequences with h-region lengths equal and longer, or less than 16 amino acids in yeast and humans.

However, an analysis of h-region length distributions supports a species-specific bias. Only 11% (33 of 300) of annotated *S. cerevisiae* signal peptides contain h-regions of length 16 or more residues, compared to 21.4% (732 of 3,415) for human signal peptides (Fig. 5C). While the total number of annotated yeast signal peptides is likely underestimated—considering that ∼30% of the ∼6,000 proteins in the yeast proteome are predicted to transit the ER—the available database trends confirm that *S. cerevisiae* signal peptides tend to have shorter h-regions than their human counterparts.

### Membrane thinning of the SPC TM window in yeast and human SPC

To understand the molecular basis of the h-region length difference between yeast and human signal peptides, we used coarse-grained molecular dynamics (MD) simulations to compare membrane thinning induced by the yeast SPC (AlphaFold-predicted structure) and the human SPC (cryo-EM structure), embedded in their respective ER membrane models.

Although the predicted yeast and cryo-EM human structures show a high degree of structural similarity, MD simulations revealed that the yeast SPC induces more membrane thinning within the TM window than the human complex (Fig. 6). The bulk yeast ER membrane was slightly thinner than the human ER membrane (38.1 versus 40.1 Å), consistent with their different lipid compositions. However, the difference was greater within the TM window: the thickness of the yeast TM window was estimated at ∼20 Å, whereas the human TM window measured ∼25 Å (Fig. 6A). Relative to their respective bulk membranes, this corresponded to 47.5% thinning for the yeast SPC and 38.3% for the human SPC (Fig. 6B). This 5 Å difference corresponds to approximately three residues in an α-helix, providing a molecular explanation for how longer human signal peptides can bind the thicker human TM window and be cleaved by the human SPC but are excluded from the thinner TM window of the yeast SPC and thus remain uncleaved.

**Figure 6.**
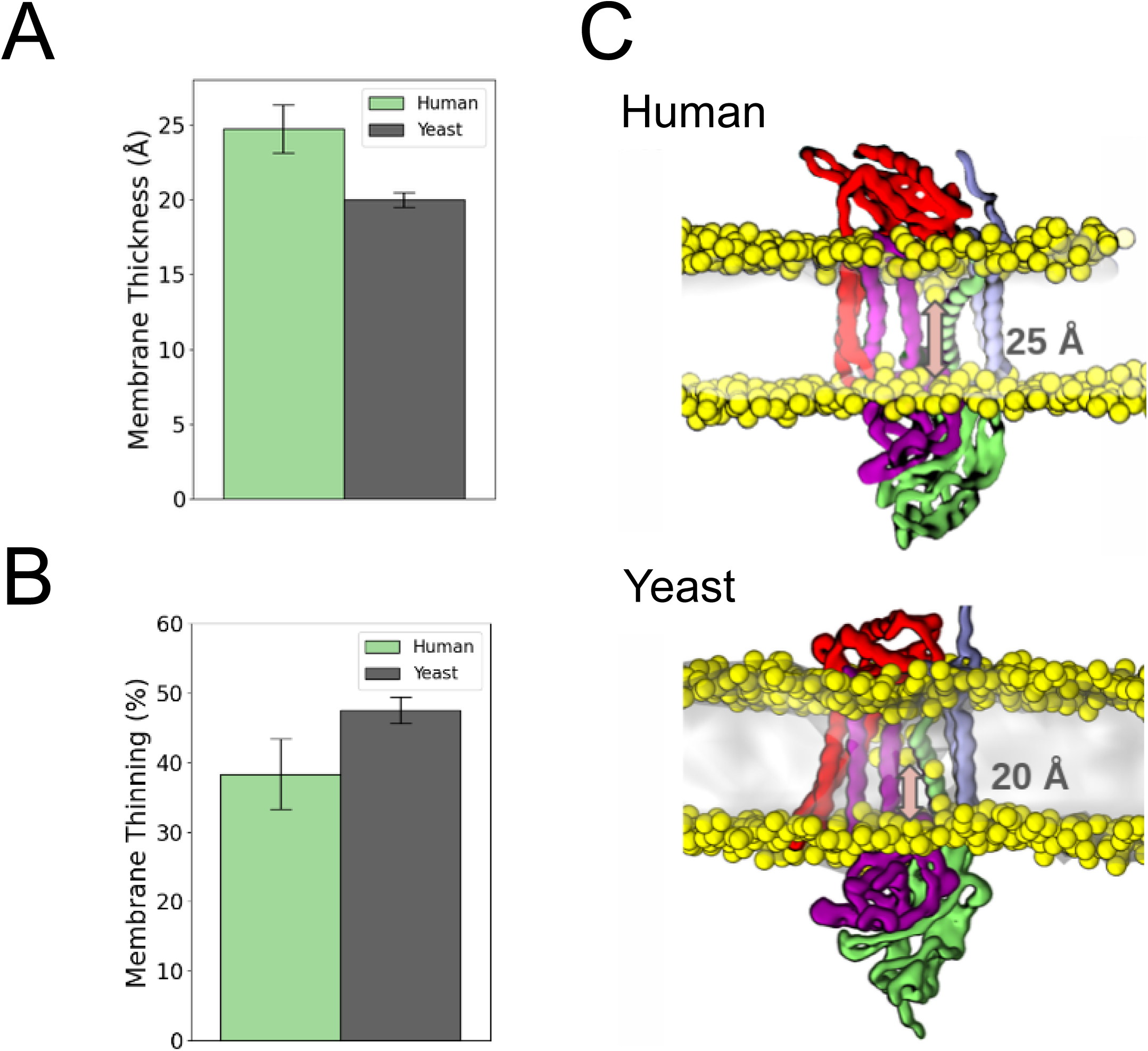
Species-specific membrane thickness and thinning within the TM window of human and yeast SPCs. (A) Average membrane thickness within the TM window of the human and yeast SPCs, calculated from the distance between the phosphate-density peaks of the two membrane leaflets. (B) Membrane thinning within the TM window relative to the corresponding protein-free bulk ER membrane. Bars represent the mean across four independent simulation replicas, and error bars indicate the standard deviation. (C) Representative simulation snapshots of the human and yeast SPCs embedded in their respective ER membrane models. Lipid phosphate groups are shown as yellow spheres. Arrows indicate the average distance between the two phosphate layers within the TM window, corresponding to approximately 25 Å for the human SPC and 20 Å for the yeast SPC.

## Discussion

Despite the high conservation of the protein secretion machinery between yeast and mammals, not all signal peptides function interchangeably between them (23–25). Previous studies attributed these incompatibilities primarily to differences in SRP recognition and translocation efficiency (26, 43). Our work demonstrates that signal peptide cleavage by the SPC constitutes an additional, previously underappreciated determinant of species-specific compatibility.

Our experimental data show that human signal peptides with longer h-regions are properly targeted and translocated across the yeast ER membrane. However, the majority escaped cleavage by the yeast SPC and consequently remained as signal-anchor sequences. These observations identify SPC-mediated processing as a main source of incompatibility for this class of human signal peptides in yeast.

Our analysis also determined that the 50% cleavage threshold for signal sequences is ∼3 residues shorter for yeast SPC compared to that for human SPC. Interestingly, this difference corresponds to approximately one turn of an alpha helix (∼4.5 Å), which closely matches the membrane thickness difference between yeast and human SPC TM windows estimated by MD simulations (∼5 Å). Together, these observations support a model in which the membrane-thinned TM window of the SPC functions as a species-specific molecular ruler. Signal peptides whose hydrophobic cores match the thickness of the TM window can adopt a cleavage-competent conformation, whereas longer hydrophobic segments may be poorly accommodated within the thinner yeast SPC TM window. This model is further supported by the observation that endogenous signal peptides of *S. cerevisiae* tend to be shorter than those of humans.

One unexpected finding from our MD simulations was that the yeast and human SPCs induced different degrees of membrane thinning despite their similar overall architectures. Because these simulations were originally performed in separate studies and analyzed using different procedures, we re-analyzed both datasets using the same method. Although the species-specific membrane models differed slightly in their bulk thicknesses because of their distinct lipid compositions, this difference alone cannot account for the approximately 5 Å difference observed within the TM window. Relative to their respective bulk membranes, the yeast SPC induced greater thinning than the human SPC, although the human system exhibited larger local fluctuations and consequently greater variability in the calculated thickness. The smaller bulk thickness of the yeast membrane was expected given its enrichment in shorter-chain lipids, particularly DYPC (1,2-dipalmitoleoyl-*sn*-glycero-3-phosphocholine) and YOPC (1-palmitoleoyl-2-oleoyl-*sn*-glycero-3-phosphocholine). However, the greater local thinning within the yeast SPC TM window may also reflect differences in the residues lining this region. The yeast complex contains polar residues that can interact with lipid headgroups and promote their displacement toward the membrane interior. Consistent with this interpretation, mutation of these residues was previously shown to increase membrane thickness within the TM window (18). Together, these observations suggest that species-specific thinning arises from the combined effects of membrane composition and the amino acid composition of the SPC TM window.

Our findings also have practical implications for recombinant protein production in yeast, where the choice of signal peptide is often a key determinant of secretion efficiency. Although numerous signal peptides have been evaluated empirically, their performance remains difficult to predict because the molecular basis of species-specific compatibility has remained poorly understood. Our results suggest that engineering the h-region length to match the cleavage preference of the host SPC may provide a rational strategy for improving secretion efficiency.

Together, our biochemical, bioinformatic, and computational analyses support a model in which the membrane-thinned TM window of the SPC functions as a molecular ruler that defines species-specific h-region length thresholds for signal peptide cleavage. More broadly, our findings suggest that substrate selectivity is determined not solely by the SPC architecture, but also by the local membrane environment created through SPC-lipid interactions. This work provides a mechanistic framework for understanding both the evolution of signal peptide processing and the rational engineering of recombinant protein secretion.

## Materials and Methods

### Yeast strains

The *Saccharomyces cerevisiae* haploid W303-1α (*MATα*, *ade2*, *can1*, *his3*, *leu2*, *trp1*, *ura3*) was used as a WT strain. *spc3-*4 is a temperature-sensitive mutant, exhibiting a defect in the SPC activity at 37°C (44).

### Construction of plasmids

Construction of LepCC variants: LepCC variants (14L, 17L, 20L, and 23L) generated in a previous study (31) were used as templates for site-directed mutagenesis to generate the 15L, 16L, 18L, and 19L variants using the KOD Mutagenesis Kit (Toyobo). All LepCC variants were cloned into the pRS424GPD yeast expression vector and fused to a C-terminal 3×HA epitope tag. The resulting yeast expression constructs were subsequently transferred into the pcDNA5-FRT/TO mammalian expression vector by Gibson assembly.

Construction of human signal peptide fusion proteins: cDNA templates for amplification of human genes were synthesized from mRNA isolated from HEK293T, HeLa, HAP1, and HepG2 cells using the SuperScript™ IV First-Strand Synthesis System (Invitrogen). DNA fragments encoding the N-terminal 60 amino acids of ADA21, AGRG2, BT2A2, CD320, CNPY3, HIPL2, NMU, PCD16, PLXA1, and RAE1E (65 amino acids for ATS4) were amplified by PCR from the corresponding cDNA templates and subcloned into yeast and mammalian expression vectors containing signal sequence-deficient CPYt (residues from 36 to 323 of CPY) fused to a C-terminal 3×HA epitope tag using Gibson assembly. Variants of ADA21, NMU, and PCD16 containing two additional leucine residues were generated by site-directed mutagenesis.

Construction of yeast proteins: DNA fragments encoding full-length or truncated Atg15, Gda1, Hoc1, Ktr3, Mnn10, Mnn11, and Sec11 were amplified by PCR and subcloned into the pRS424GPD yeast expression vector and the pcDNA5-FRT/TO mammalian expression vector with a C-terminal 3×HA epitope tag using Gibson assembly. Two C-terminal N-linked glycosylation sites were introduced by site-directed mutagenesis.

### Mammalian cell culture

Flp-In T-REx 293 cells (Invitrogen; R78007) transfected with pSpCas9(BB)-2A-Puro (PX459) V2.0 containing non-targeting sgRNA were used (a kind gift from Professor Sang Wook Kang in University of Ulsan College of Medicine, South Korea). Cells were grown in Dulbecco’s Modified Eagle’s Medium (DMEM) supplemented with 10% fetal calf serum and 0.5 μg/mL puromycin, and incubated in 5% CO_2_ at 37°C. The Flp-In T-REx 293-Cas9 cells were transfected with pcDNA5-FRT/TO vector bearing protein of interest and Lipofectamine® 3000 reagent (Invitrogen). After 7 hr of transfection, protein expression was induced by the treatment with doxycycline (10 ng/mL). For the cavinafungin treatment, cells were treated with 10 uM cavinafungin for 14 hr after 1 hr of induction.

### Cell lysis and SDS-PAGE

Yeast cells were grown overnight in selective medium lacking tryptophan (-Trp) to the logarithmic growth phase. Cells corresponding to 1 OD_600_ unit were harvested and stored at −20°C until use. Cell pellets were resuspended in 500 μL of distilled water and mixed with alkaline lysis buffer (1.85 N NaOH, 10 mM PMSF, and 1.05 M β-mercaptoethanol). Samples were vigorously vortexed and incubated on ice for 10 min. Trichloroacetic acid (TCA; 50%, 575 μL) was added to achieve a final concentration of 25%, followed by incubation on ice for 1 h. Protein precipitates were collected by centrifugation at 20,238 × *g* and 4°C for 30 min. The pellets were washed with 100% acetone and resuspended in 50 μL of sample buffer containing 50 mM DTT, 50 mM Tris-HCl (pH 7.6), 5% SDS, 5% glycerol, 50 mM EDTA, 1 mM PMSF, and 1× protease inhibitor cocktail (Quartett). Samples were incubated at 30°C for 10 min with vigorous mixing and centrifuged at 20,238 × *g* for 2 min at room temperature. The supernatants were transferred to fresh tubes and treated with Endo Hf (New England Biolabs) at 37°C for 1 h before separation by SDS-PAGE.

Mammalian cells cultured in 12-well plates were lysed in 500 μL of lysis buffer (1% SDS, 100 mM Tris-HCl [pH 7.6], 1 mM PMSF, and 1× protease inhibitor cocktail). Cell lysates were boiled at 95°C for 3 min and vortexed; this process was repeated three times until the lysates were no longer viscous. Lysates were centrifuged at 20,238 × *g* for 30 s at room temperature and treated with either Endo Hf or PNGase F (Enzynomics) at 37°C for 1 h. Protein samples were then mixed with 5× Laemmli sample buffer, boiled at 95°C for 5 min, and subjected to SDS-PAGE.

### Western blotting

Proteins were transferred onto nitrocellulose membranes using a wet-transfer system. Membranes were blocked with 5% skim milk in TBS and incubated with mouse anti-HA primary antibody (1:10,000; BioLegend) diluted in TBST. After three washes with TBST, membranes were incubated with HRP-conjugated goat anti-mouse secondary antibody (1:50,000; Rockland) diluted in TBST. Following three additional washes with TBST, immunoreactive bands were detected using ChemiDoc^TM^ XRS+.

### Data quantification

Cell growth assay and blots were detected by ChemiDoc^TM^ XRS+ and resulting data were processed using Image Lab^TM^ software.

### Statistical analysis

Statistical analyses with obtained quantification data were performed using MATLAB.

### Analysis of *S. cerevisiae* and *H. sapiens* signal peptide lengths

Signal peptides of *S. cerevisiae* and *H. sapiens* were downloaded from Signal peptide explorer (https://services.healthtech.dtu.dk/services/SPEx-1.0/). n-, h- and c-regions of signal peptides were sorted using SignalP 6.0 slow mode (39). Full-length protein sequences were used as input for SignalP 6.0 predictions.

### Coarse-grained MD-simulations

Previously generated coarse-grained molecular dynamics trajectories of the apo human and yeast signal peptidase complexes (SPCs), embedded in their respective endoplasmic-reticulum membrane models, were reused for this analysis. The construction of the systems and the simulation protocols have been described previously for the human (17) and yeast (18) SPCs. Briefly, the simulations were performed using the Martini 3 force field and comprised five independent 20-µs replicas for each complex. The human and yeast signal peptidase complexes (SPCs) were embedded into symmetric endoplasmic reticulum (ER) membrane models for each species. The human ER membrane was composed of a POPC:POPE:POPS:Cholesterol:PI(3,4)P2 mixture at a molar ratio of 44:26:4:15:11 whereas the yeast ER Membrane was constructed according to the Monje-Galvan & Klauda (2015) ER model (45), composed of DYPC:YOPC:POPI:PYPI:DYPE:YOPE:ERGO:YOPA:YOPS:POPS at a ratio of 42:28:21:14:10:10:7:6:6:6. Additional details regarding system construction and simulation protocols are provided in the original publications(17, 18).

Because previous studies evaluated human and yeast SPC simulations using different analysis routines, all production trajectories were re-analyzed here using a single unified protocol to ensure consistency. To eliminate edge artifacts, a trajectory in the human ER simulations dataset was omitted because lipid PO_4_ headgroups in the lower leaflet were displaced outside the spatial selection threshold centered around the protein transmembrane (TM) window. To maintain balanced sampling across species for subsequent analysis, the simulation representing the lowest thickness in the yeast dataset was also discarded. Four production trajectories per species were retained for analysis. The PO_4_ density profiles from the SPC-containing simulations were fitted using a triple-Gaussian function, whereas those from the corresponding protein-free reference membranes were fitted using a double-Gaussian function.

The local membrane thickness, d_TM_ window, was obtained directly from the separation between the fitted phosphate-density peaks in the TM-window region. The bulk membrane thickness, d_bulk,_ was determined analogously from the corresponding protein-free reference membrane. Membrane thinning was then calculated relative to the bulk membrane thickness as:

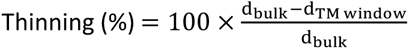

## Acknowledgments

Authors thank Professor Sang Wook Kang (University of Ulsan College of Medicine, South Korea) for a kind gift of HEK293T cell-lines and Sungjoon Son for technical assistance. This work was supported by the National Research Foundation of Korea(NRF) grant funded by the Korea government (MSIT)(RS-2023-00259499 and RS-2025-00564001) to HK. Computer hours awarded to the GENCI project numbers 2022-A0120713456 and 2023-A014071345 on the Jean-Zay (IDRIS), Adastra (CINES) and Joliot-Curie (TGCC) clusters of the French National Supercomputing Center and the computing time available at the IN2P3 computer cluster in Lyon are gratefully acknowledged. P.C.T. Souza acknowledge the support provided by the CNRS and Sanofi. P.C.T. Souza and G. P. Pereira also acknowledge the support of the Centre Blaise Pascal’s IT test platform at ENS de Lyon (Lyon, France) for the computer facilities. The platform operates the SIDUS solution developed by Emmanuel Quemener (46). G.P.P also gratefully acknowledges the support from the Torres Quevedo grant PTQ2023-012988, financed by the MCIU/AEI/10.13039/501100011033, and from Zymvol Biomodeling S.L.

## Author Contributions

Y. Chung and H. Kim, experimental conceptualization; G. P. Pereira, and P.C.T. Souza, computational conceptualization. Y. Chung, G. P. Pereira, Y. J. Lee, L. R. Kjølbye, P.C.T. Souza, data curation, formal analysis, investigation, validation, visualization; P.C.T. Souza and H. Kim, funding acquisition, project administration, supervision; Y. Chung, G. P. Pereira, P.C.T. Souza and H. Kim, writing-original draft; Y. Chung, G. P. Pereira, Y. J. Lee, A. Agnesa, L. R. Kjølbye, H. Nielsen, G. von Heijne, P.C.T. Souza and H. Kim, writing-review & editing.

## Conflict of interest

Authors declare no conflict of interest.

---

**Supplemental Figure S1.**
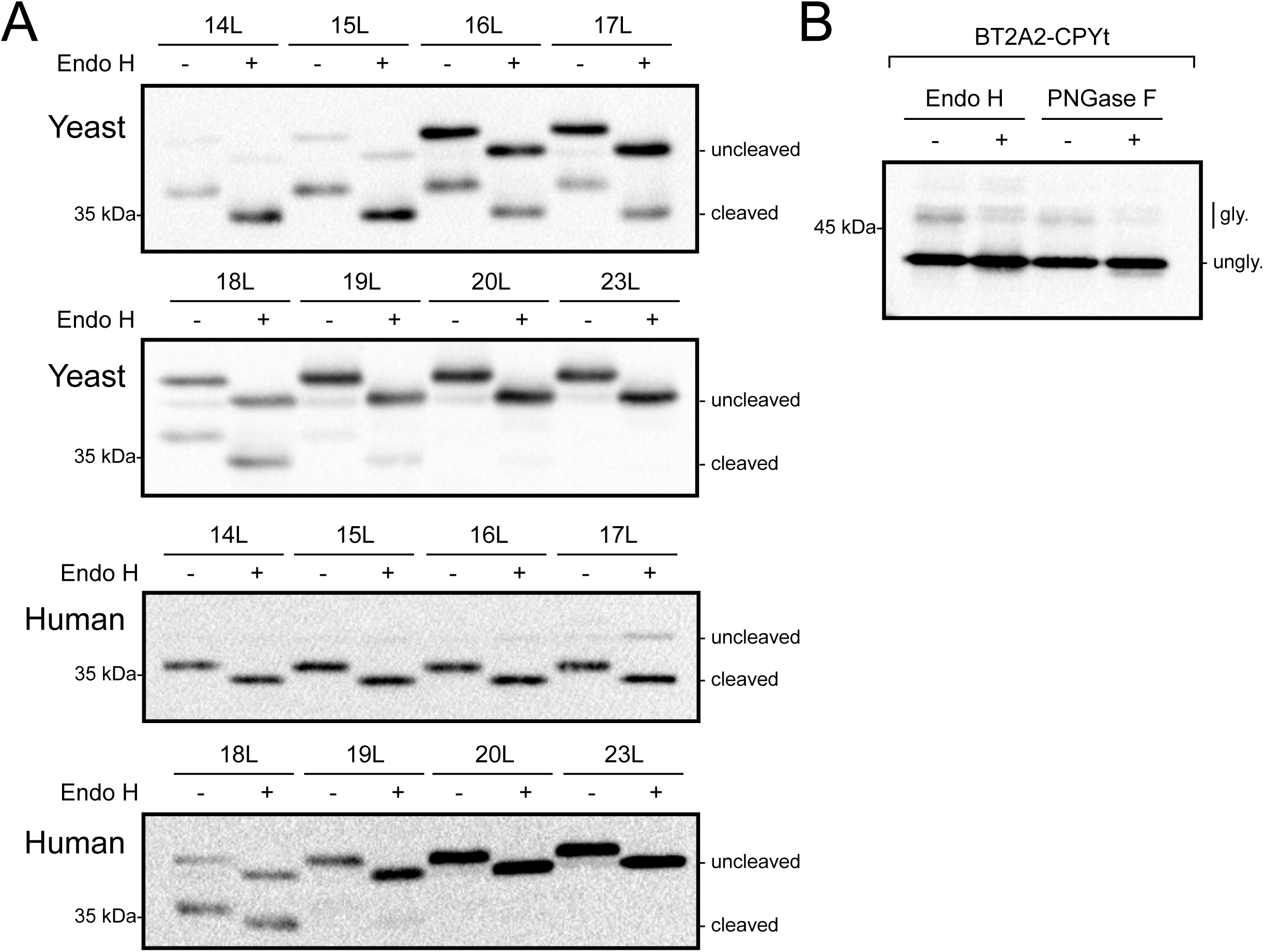
Comparison of SPC-mediated cleavage of LepCC variants in yeast and human cells. (A) Western blot analysis of LepCC variants expressed in yeast and human cells with or without endoglycosidase H (Endo H) treatment. (B) Western blot analysis of the BT2A2 fusion protein expressed in HEK293 cells following treatment with Endo H or PNGase F.

**Supplemental Figure S2.**
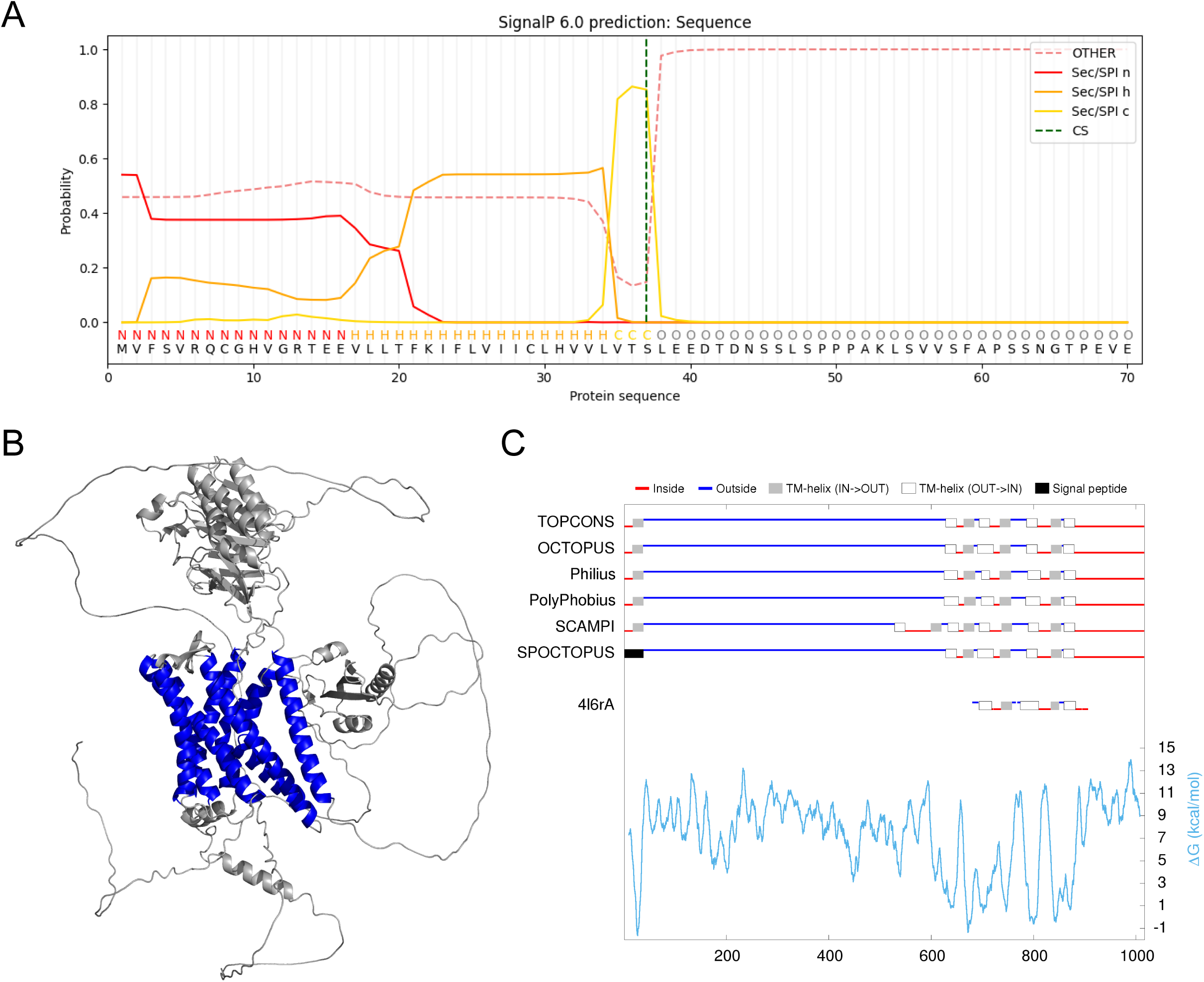
Predicted structural features and membrane topology of AGRG2. (A) SignalP 6.0 (39)-predicted n-, h-, and c-regions of AGRG2. (B) AlphaFold-predicted structure of AGRG2. Predicted transmembrane (TM) domains are shown in blue. (C) TOPCONS (47)-predicted signal peptide (black) and TM helices (gray) of AGRG2.

**Supplemental Figure S3.**
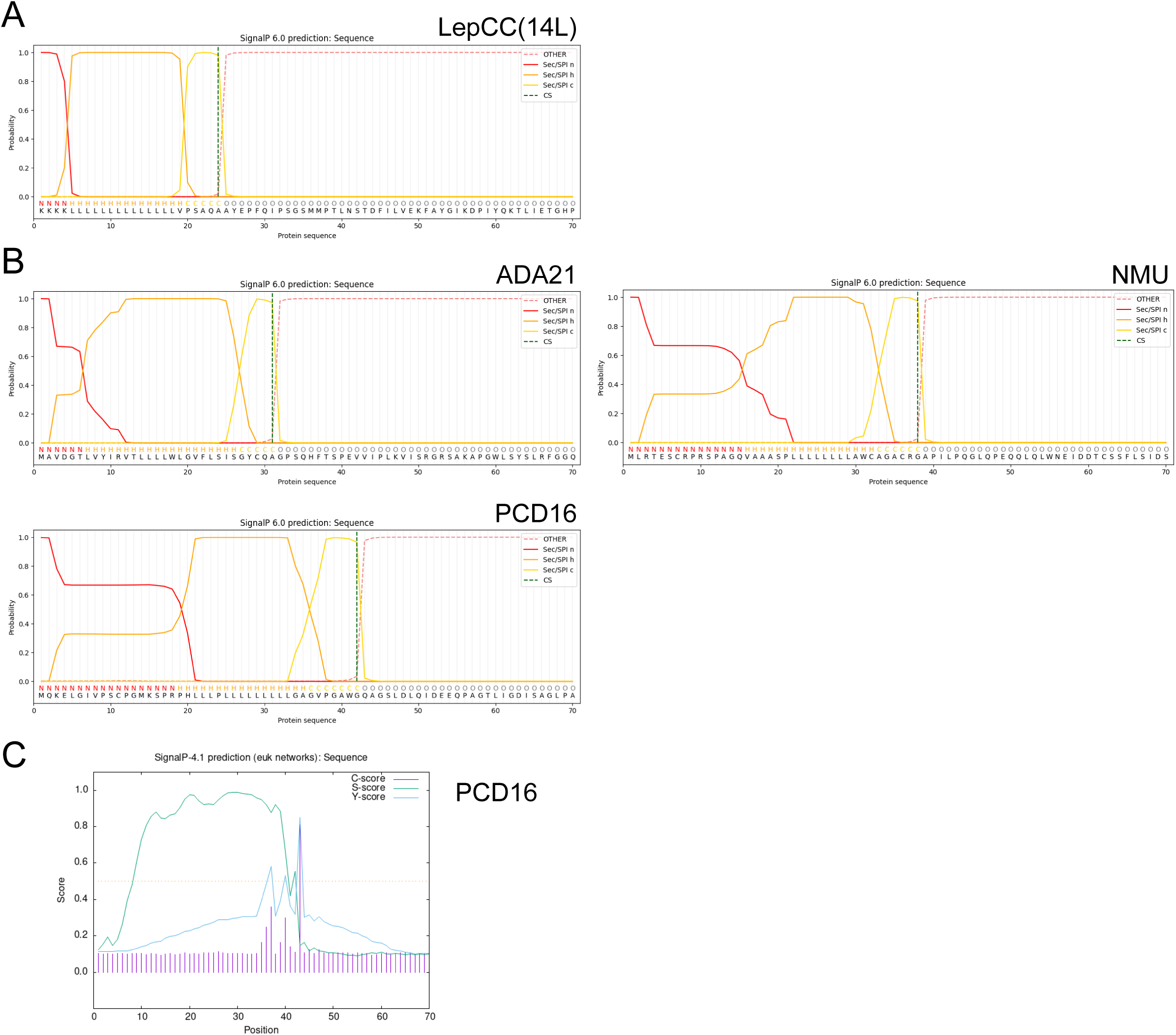
SignalP-predicted signal peptide regions and cleavage sites. SignalP 6.0 (39)-predicted n-, h-, and c-regions and cleavage sites of (A) TM2 of LepCC(14L) and (B) the signal peptides of ADA21, NMU, and PCD16. (C) SignalP 4.1 (40)-predicted cleavage sites of PCD16.

**Supplemental Figure S4.**
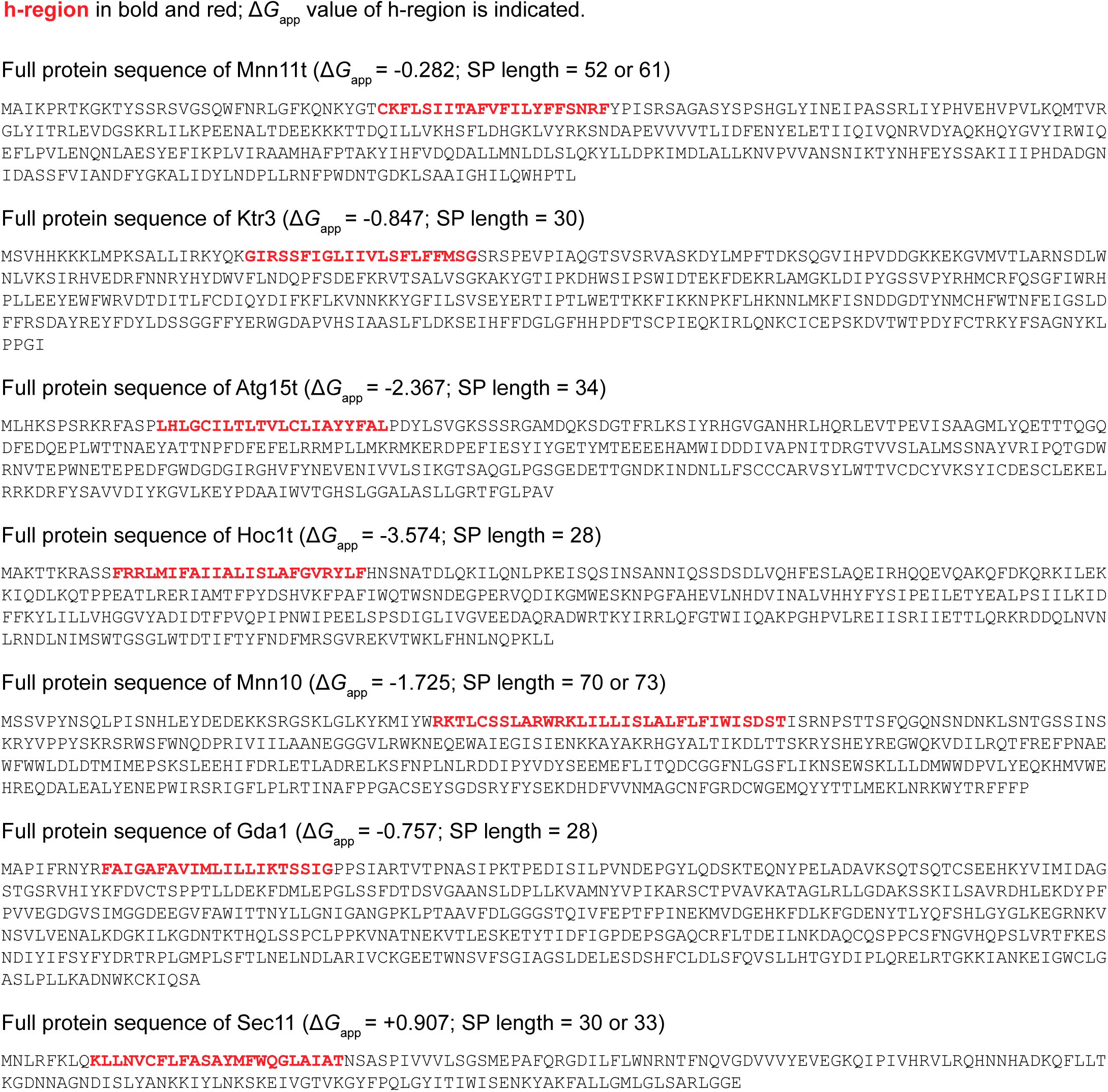
Sequences of yeast signal-anchor proteins used in this study.

## Notes

### Competing Interest Statement

The authors have declared no competing interest.

